# The hippocampus as a predictive map

**DOI:** 10.1101/097170

**Authors:** Kimberly L. Stachenfeld, Matthew M. Botvinick, Samuel J. Gershman

## Abstract

A cognitive map has long been the dominant metaphor for hippocampal function, embracing the idea that place cells encode a geometric representation of space. However, evidence for predictive coding, reward sensitivity, and policy dependence in place cells suggests that the representation is not purely spatial. We approach this puzzle from a reinforcement learning perspective: what kind of spatial representation is most useful for maximizing future reward? We show that the answer takes the form of a predictive representation. This representation captures many aspects of place cell responses that fall outside the traditional view of a cognitive map. Furthermore, we argue that entorhinal grid cells encode a low-dimensional basis set for the predictive representation, useful for suppressing noise in predictions and extracting multiscale structure for hierarchical planning.

## Introduction

Learning to predict long-term reward is fundamental to the survival of many animals. Some species may go days, weeks or even months before attaining primary reward, during which time aversive states must be endured. Evidence suggests that the brain has evolved multiple solutions to this reinforcement learning (RL) problem^1^. One solution is to learn a model or “cognitive map” of the environment^2^, which can then be used to generate long-term reward predictions through simulation of future states^1^. However, this solution is computationally intensive, especially in real-world environments where the space of future possibilities is virtually infinite. An alternative “model-free” solution is to learn, from trial-and-error, a value function mapping states to long-term reward predictions^3^. However, dynamic environments can be problematic for this approach, because changes in the distribution of rewards necessitates complete relearning of the value function.

Here, we argue that the hippocampus supports a third solution: learning of a “predictive map” that represents each state in terms of its “successor states” (upcoming states)^4,5^. This representation is sufficient for long-term reward prediction, is learnable using a simple, biologically plausible algorithm, and explains a wealth of data from studies of the hippocampus.

Our primary focus is on understanding the computational function of hippocampal place cells, which respond selectively when an animal occupies a particular location in space^6^. A classic and still influential view of place cells is that they collectively furnish an explicit map of space^7,8^. This map can then be employed as the input to a model-based^9–11^ or model-free^12,13^ RL system for computing the value of the animal's current state. In contrast, the predictive map theory views place cells as encoding predictions of future states, which can then be combined with reward predictions to compute values. This theory can account for why the firing of place cells is modulated by variables like obstacles, environment topology, and direction of travel. It also generalizes to hippocampal coding in non-spatial tasks. Beyond the hippocampus, we argue that entorhinal grid cells^14^, which fire periodically over space, encode a low-dimensional decomposition of the predictive map, useful for stabilizing the map and discovering subgoals.

## Results

### The successor representation

An animal's optimal course of action will frequently depend on the location (or more generally, the “state”) that the animal is in. The hippocampus' purported role of representing location is therefore considered to be a very important one. The traditional view of state representation in the hippocampus is that the place cells index the current location by firing when the animal visits the encoded location, remaining silent otherwise^7^. The main idea of the SR model, elaborated below, is that place cells do not encode place *per se*, but rather a predictive representation of future states given the current state. Two states that predict similar future states will have similar representations, and two physically adjacent states that predict divergent future states will have dissimilar representations.

To motivate our use of the SR in the RL setting, we demonstrate that this representation emerges naturally as a term *M* in the definition of value (*V*) often used in RL. We consider the problem of RL in a Markov decision process consisting of the following elements^15^: a set of states (e.g., spatial locations), a set of actions, a transition distribution *P*(*s′|s, a*) specifying the probability of transitioning to state *s*′ from state *s* after taking action *a*, a reward function *R*(*s*) specifying the expected immediate reward in state *s*, and a discount factor *γ ∊* [0, 1] that down-weights distal rewards. An agent chooses actions according to a policy *π*(*a|s*) and collects rewards as it moves through the state space. The value of a state is defined formally as the expected discounted cumulative future reward under policy *π*:

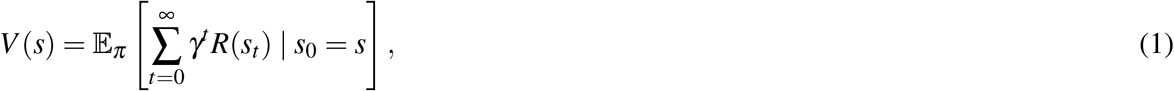

where *s_t_* is the state visited at time *t*. Our focus here is on policy evaluation (computing *V*). In our simulations we feed the agent the optimal policy; in the Supplemental Methods we discuss algorithms for policy improvement. To simplify notation, we assume implicit dependence on *π* and define the state transition matrix *T*, where *T* (*s, s*′) = ∑*_a_ π*(*a|s*)*P*(*s′|s, a*).

The value function can be decomposed into the inner product of the reward function with a predictive state representation known as the successor representation (SR)^4^, denoted by *M*:

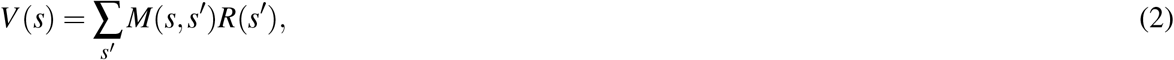

The SR encodes the expected discounted future occupancy of state *s*′ along a trajectory initiated in state *s*:

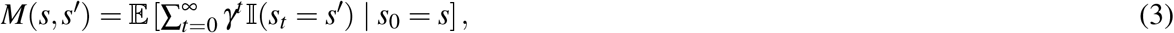

where 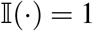 if its argument is true, and 0 otherwise.

An estimate of the SR (denoted *M̂*) can be incrementally updated using a form of the temporal difference learning algorithm^4,16^. After observing a transition *s_t_ → s_t_*_+1_, the estimate is updated according to:

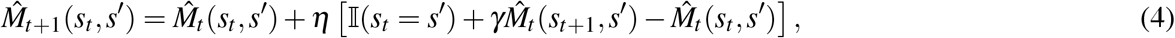

where *η* is a learning rate (unless specified otherwise, *η* = 0.1 in our simulations). The form of this update is identical to the temporal difference learning rule for value functions^15^, except that in this case the reward prediction error is replaced by a *successor prediction error* (the term in brackets). Note that these prediction errors are distinct from state prediction errors used to update an estimate of the transition function^17^; the SR predicts not just the next state but a superposition of future states over a possibly infinite horizon. The transition and SR functions only coincide when *γ* = 0. We assume the SR matrix *M* is initialized to the identity matrix, meaning *M*(*s, s*′) = 1 if *s* = *s*′, and *M*(*s, s*′) = 0 if *s* ≠ *s*′. This initialization can be understood to mean that each state will necessarily predict only itself.

The SR combines some of the advantages of model-free and model-based algorithms. Like model-free algorithms, policy evaluation is computationally efficient with the SR. However, factoring the value function into a state dynamics SR term and a reward term confers some of the flexibility usually associated with model-based methods. Having separate terms for state dynamics and reward permits rapid recomputation of new value functions when reward is changed independently of state dynamics, as demonstrated in Fig. 1. The SR can be learned before any reward has been seen, so that at the first introduction of reward, a value function can be computed immediately. When the reward function changes – such as when the animal becomes satiated, or when food is redistributed about the environment – the animal can immediately recompute a new value function based on its expected state transitions. A model-free agent would have to relearn value estimates for each location in order to make value predictions, and a model-based agent would need to aggregate the results of time-consuming searches through its model before it could produce an updated value prediction^1,4^. In Fig. S2, we demonstrate that while changing the reward function completely disrupts model free learning of a value function in a 2-step tree maze, SR learning can quickly adjust. Thus, the SR combines the efficiency of model-free control with some of the flexibility of model-based control.

**Figure 1.**
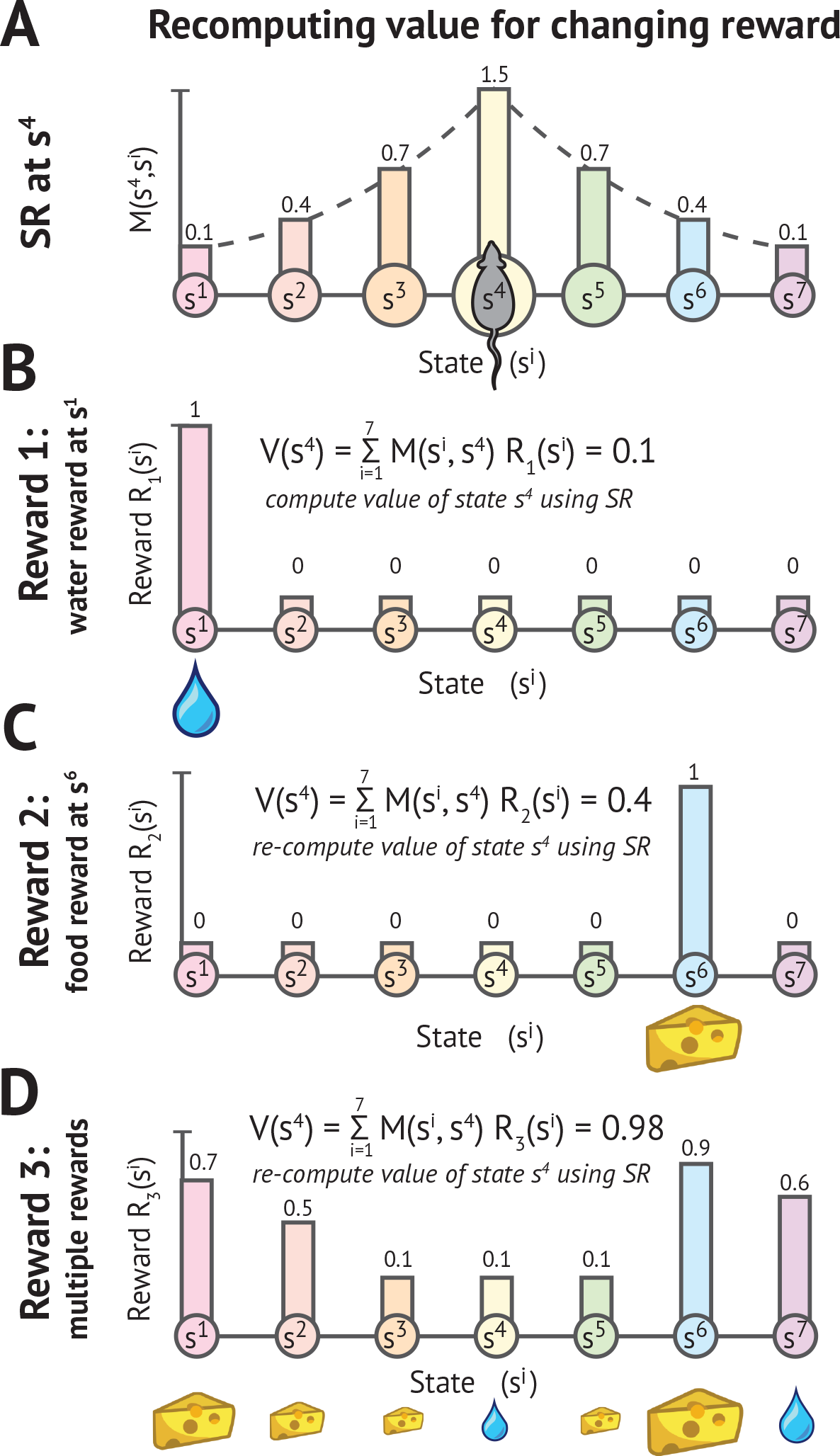
Updating value with the SR following change in reward. Since the representations of state and reward are decoupled, value functions can be rapidly recomputed for new reward functions without changing the SR. As formally defined in Equation 3, *M*(*s, s*′) gives the expected number of visits to state *s*′ given a current location of *s*. Panel A shows the successor representation of state *s*^4^, which corresponds to a row *M*(*s*^4^,:) of the SR matrix. Panels B-D show how the value of *s*_4_ changes under different reward functions.

For an agent trying to optimize expected discounted future reward, two states that predict similar successor states are necessarily similarly valuable, and can be safely grouped together^18^. This makes the SR a good metric space for generalizing value. Since adjacent states will frequently lead to each other, the SR will naturally represent adjacent states similarly and therefore be smooth over time and space in spatial tasks. Since the SR is well defined for any Markov decision process, we can use the same architecture for many kinds of tasks, not just spatial ones.

### Hippocampal encoding of the successor representation

We now turn to our main theoretical claim: that the SR is encoded by the hippocampus. This hypothesis is based on the central role of the hippocampus in representing space and context^19^, as well as its contribution to sequential decision making^20,21^. Although the SR can be applied to arbitrary state spaces, we focus on spatial domains where states index locations.

Place cells in the hippocampus have traditionally been viewed as encoding an animal's current location. In contrast, the predictive map theory views these cells as encoding an animal's *future* locations. Crucially, an animal's future locations depend on its policy, which is constrained by a variety of factors such as the environmental topology and the locations of rewards. We demonstrate that these factors shape place cell receptive field properties in a manner consistent with a predictive map.

According to our model, the hippocampus represents the SR as a rate code across the population. Each neuron represents some possible future state (e.g., spatial position) in the environment. At any current state *s*, the population will encode a row of the SR matrix, *M*(*s*,:). The firing rate of a single neuron encoding state *s* in the population is proportional to the discounted expected number of times it will be visited under the present policy given the current position *s*. An SR place field refers to the firing rate of a single SR-encoding neuron at each state in the task and corresponds to a column of the SR matrix, *M*(:, *s*′). This vector contains the expected number of times a single encoded state *s* will be visited under the current policy, starting from any state *s*. In general, we will refer to place fields simulated under our model as “SR receptive fields” or “SR place fields.” To summarize the relationship between the SR matrix *M* and simulated hippocampal cells: The firing of *all* the neurons at *one* state *s* is modeled by a *row M*(*s*,:) of the SR matrix *M*, and the firing of *one* neuron encoding *s* evaluated at *all* states is modeled by a *column M*(:, *s*′). This is illustrated in Fig. 1.

We first try to build some intuition for this idea, and how it relates to a more traditional view of place cells. In an open, 2D environment, the canonical place has a gradually decaying, roughly circular firing field. These are often modeled as approximately Gaussian. In such an environment, the SR place fields look essentially the same, with peaks of high firing surrounded by a radius of gradually reduced firing. The SR model makes this prediction because under a random walk, the animal is likely to visit its current location and nearby locations immediately, and more distant locations later. Thus, the states closer to the encoded location of an SR place cell will predict a higher expected discounted number of visits to the encoded location, and will elicit higher firing of the encoding cell.

Fig. 3 illustrates the experimental conditions in which the predictions of the SR model (Fig. 3C) depart from the predictions of two alternative models (Fig. 3A-B). As examples, we implement the three models for a 2D room containing an obstacle and for a 1D track with an established preferred direction of travel. The first alternative model is a Gaussian place field in which firing is related to the Euclidean distance from the field center (Fig. 3A), usually invoked for modeling place field activity in open spatial domains^22,23^. The second alternative model is a topologically sensitive place field in which firing is related to the average path length from the field center, where paths cannot pass through obstacles^13^ (Fig. 3A). Like the topological place fields and unlike the Gaussian place fields, the SR place fields respect obstacles in the 2D environment. Since states on opposite sides of a barrier cannot occur nearby in time, SR place fields will tend to be active on only one side of a barrier.

**Figure 2.**
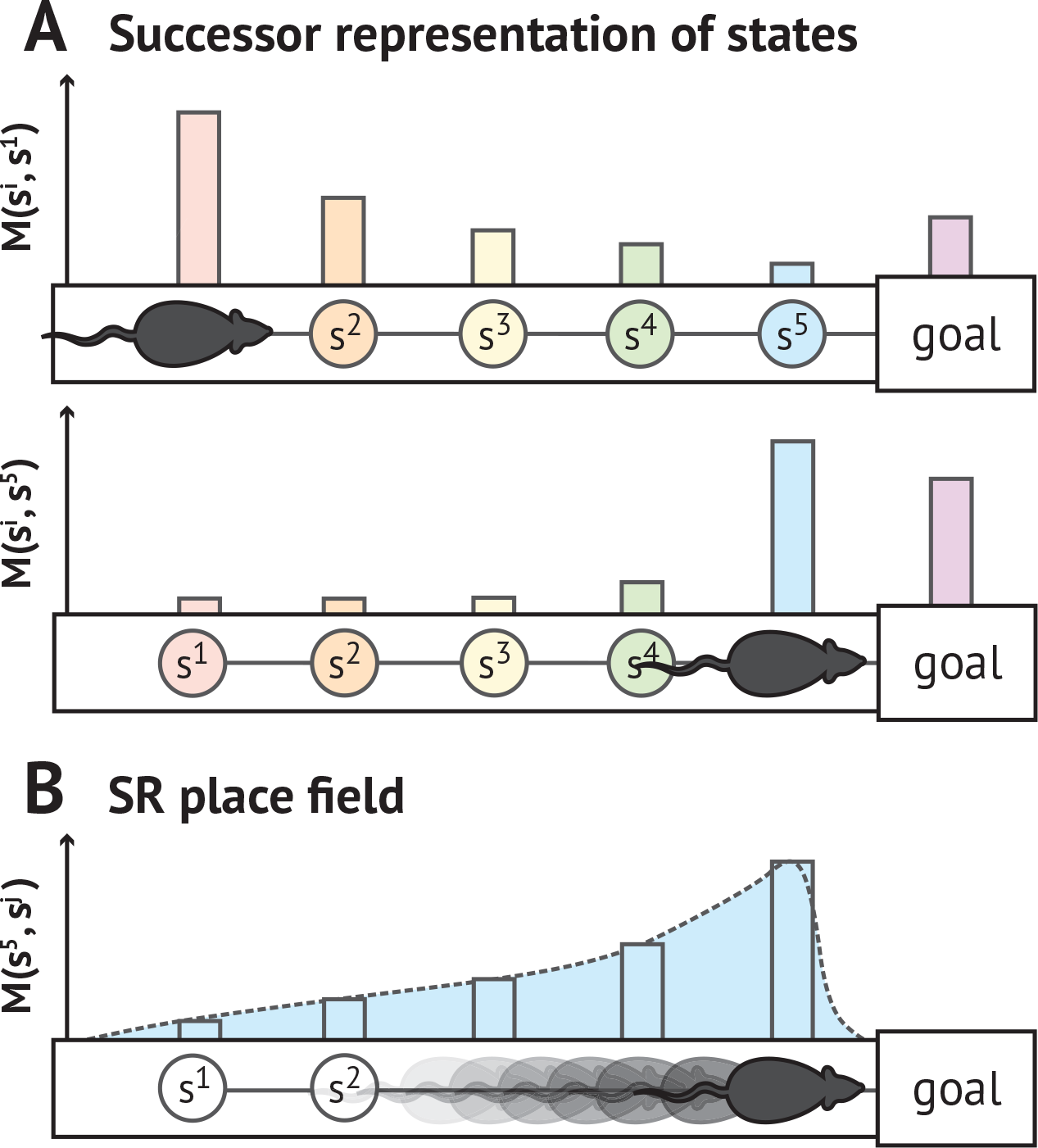
Illustration of SR encoding population and individual SR place fields. Under a prospective representation such as the SR, the population vector will be assymetrically expand in the direction of travel toward upcoming states. The place fields for individual cells will skew backwards. (A) A neural population encodes a prospective representation such that the firing rate of each cell is proportional to the discounted expected number of times its preferred state will be visited in the future. This population code is skewed toward upcoming states. Each colored bump represents the firing rate of a different place field located along the track. The value *M*(*s, s*′) is formally defined in Equation 3 as the expected number of visits to state *s*′ given a current location of *s*, and the population vectors *M*(*s*,:) illustrated here correspond to rows of the SR matrix. (B) The place field for a single SR-encoding cell skews backward toward past states that predict the cell's preferred state. When the blue state *s*^5^ is visited, it becomes associated with all past states that predicted it. This automatically assigns credit for upcoming reward to preceding states. The receptive field *M*(:, *s*′) illustrated here corresponds to a column of the SR matrix.

**Figure 3.**
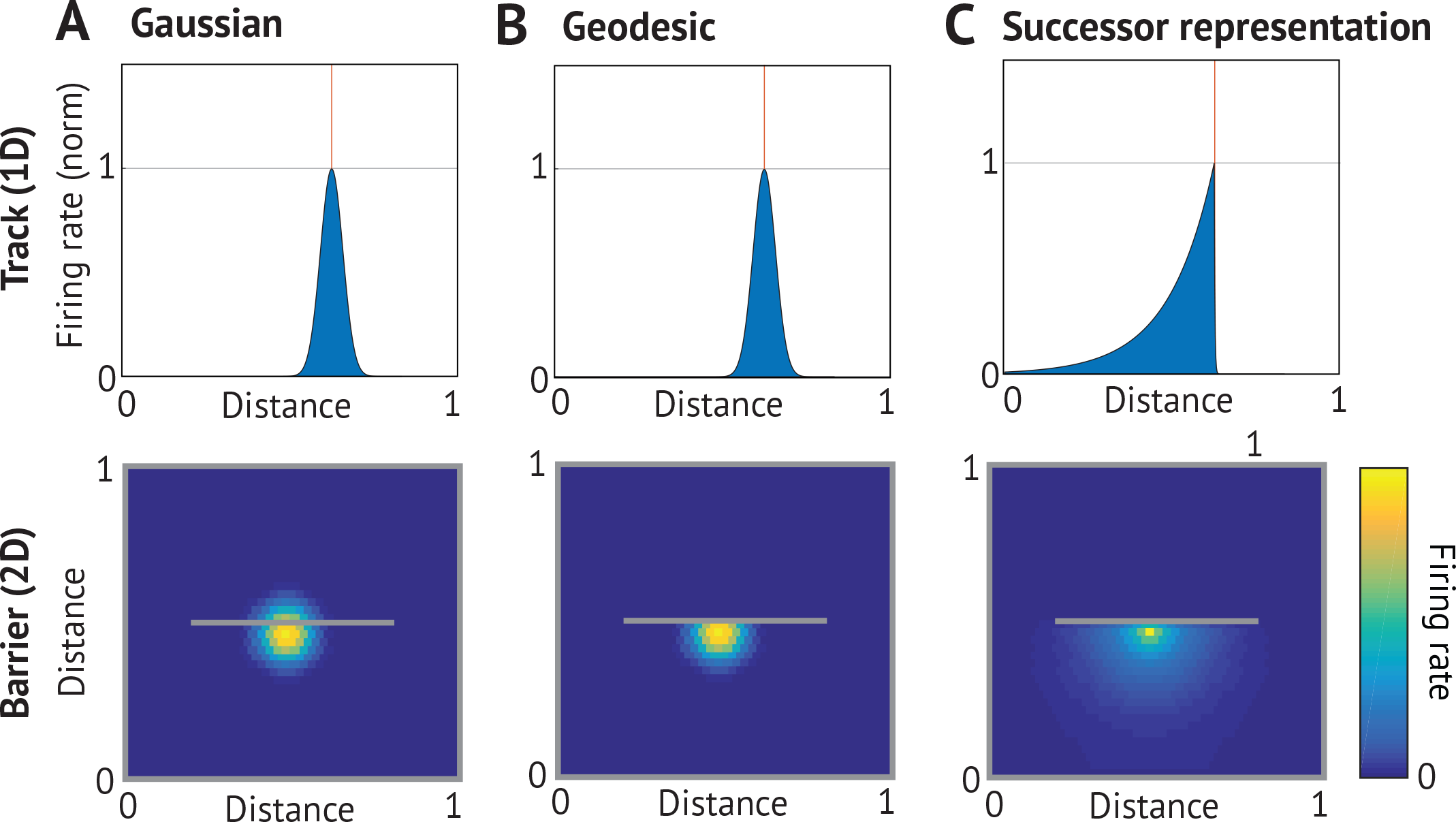
Comparison of place cell models. (Top) 1D track with left-to-right preferred direction of travel, red line marking field center; (bottom) 2D environment with a barrier indicated by gray line. (A) *Gaussian place field*. Firing of place cells decays with Euclidean distance from the center of the field regardless of experience and environmental topology. (B) *Topological place field*. Firing rate decays with geodesic distance from the center of the field, which respects boundaries in the environment but is invariant to the direction of travel^13^. (C) *SR place field*. Firing rate is proportional to the discounted expected number of visits to other states under the current policy. On the directed track, fields will skew opposite the direction of motion to anticipate the upcoming successor state. Since the policy will not permit traversing walls, successor fields warp around obstacles.

On the 1D track, the SR place fields skew opposite the direction of travel. This backward skewing arises because upcoming states can be reliably predicted further in advance when traveling repeatedly in a particular direction. Neither of the control models provide ways for a directed behavioral policy to interact with state representation, and therefore cannot show this effect. Evidence for predictive skewing comes from experiments in which animals traveled repeatedly in a particular direction along a linear track^24,25^. The authors noted this as evidence for predictive coding in hippocampus^24,26^. In Fig. 2, we explain how a future-oriented representation evokes a forward-skewing representation in the population at any given point in time but implies that receptive fields for any individual cell should skew backwards. In order for a given cell to fire predictively, it must begin firing before its encoded state is visited, causing a backward-skewed receptive field. Figure 4 compares the predicted and experimentally observed backward skewing, demonstrating that the model captures the qualitative pattern of skewing observed when the animal has a directional bias.

**Figure 4.**
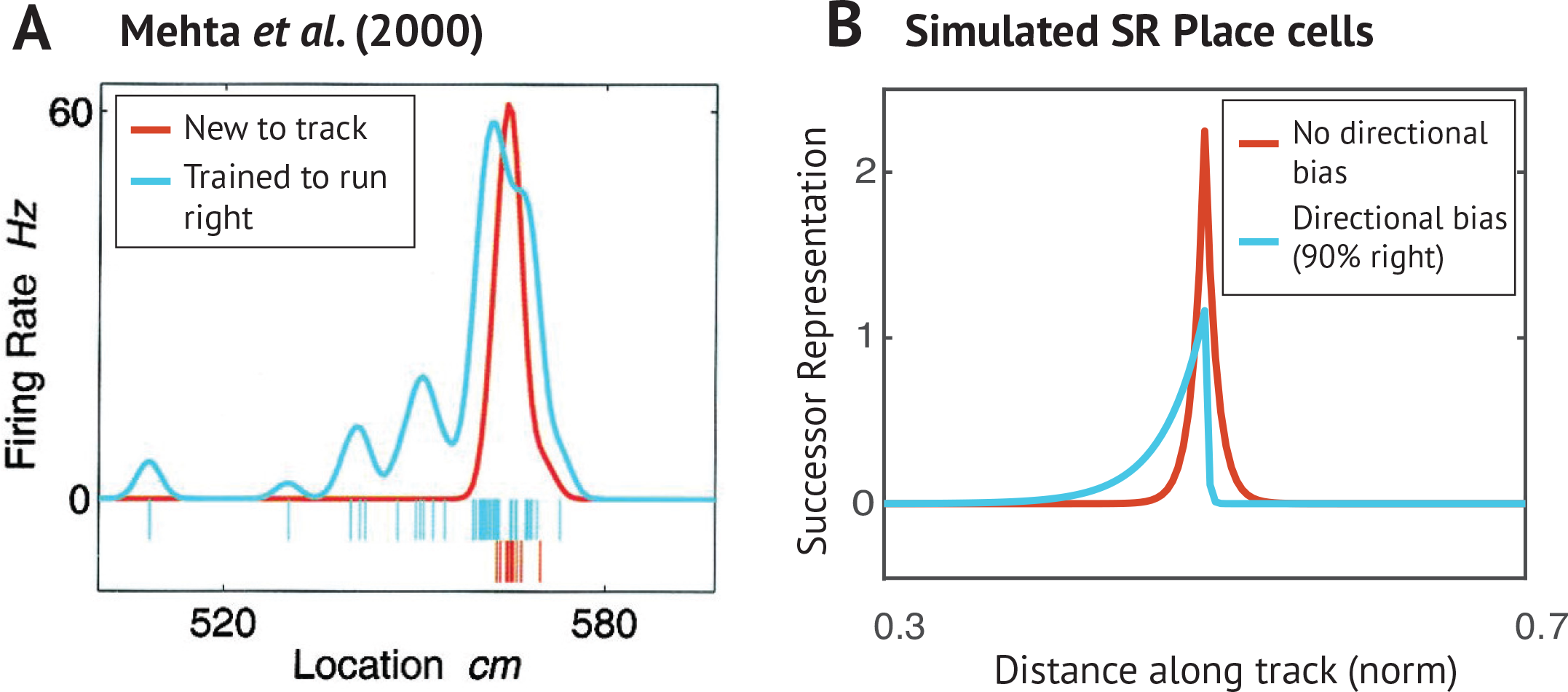
Predictive skewing of place fields. (A) As a rat is trained to run repeatedly in a preferred direction along a narrow track, initially symmetric place cells (red) begin to skew (blue) opposite the direction of travel^25^. (B) When transitions in either direction are equally probable, SR place fields are symmetric (red). Under a policy in which transitions to the right are more probable than to the left, simulated SR place fields skew opposite the direction of travel toward states predicting the preferred state (blue).

Consistent with the SR model, experiments have shown that place fields become distorted around barriers^27–29^. In Figure 5, we explore the effect of placing obstacles in a Tolman detour maze on the SR place fields and compare to experimental results obtained by Alvernhe *et al*.^29^. When a barrier is placed in a maze such that the animal is forced to take a detour, the place fields engage in “local remapping.” Place fields near the barrier change their firing fields significantly more than those further from the barrier (Fig. 5A-C). When barriers are inserted, SR place fields change their fields near the path blocked by the barrier and less so at more distal locations where the optimal policy is unaffected (Fig. 5D-F). This locality is imposed by the discount factor. The full set of place fields is included in the supplement (Fig. S3).

**Figure 5.**
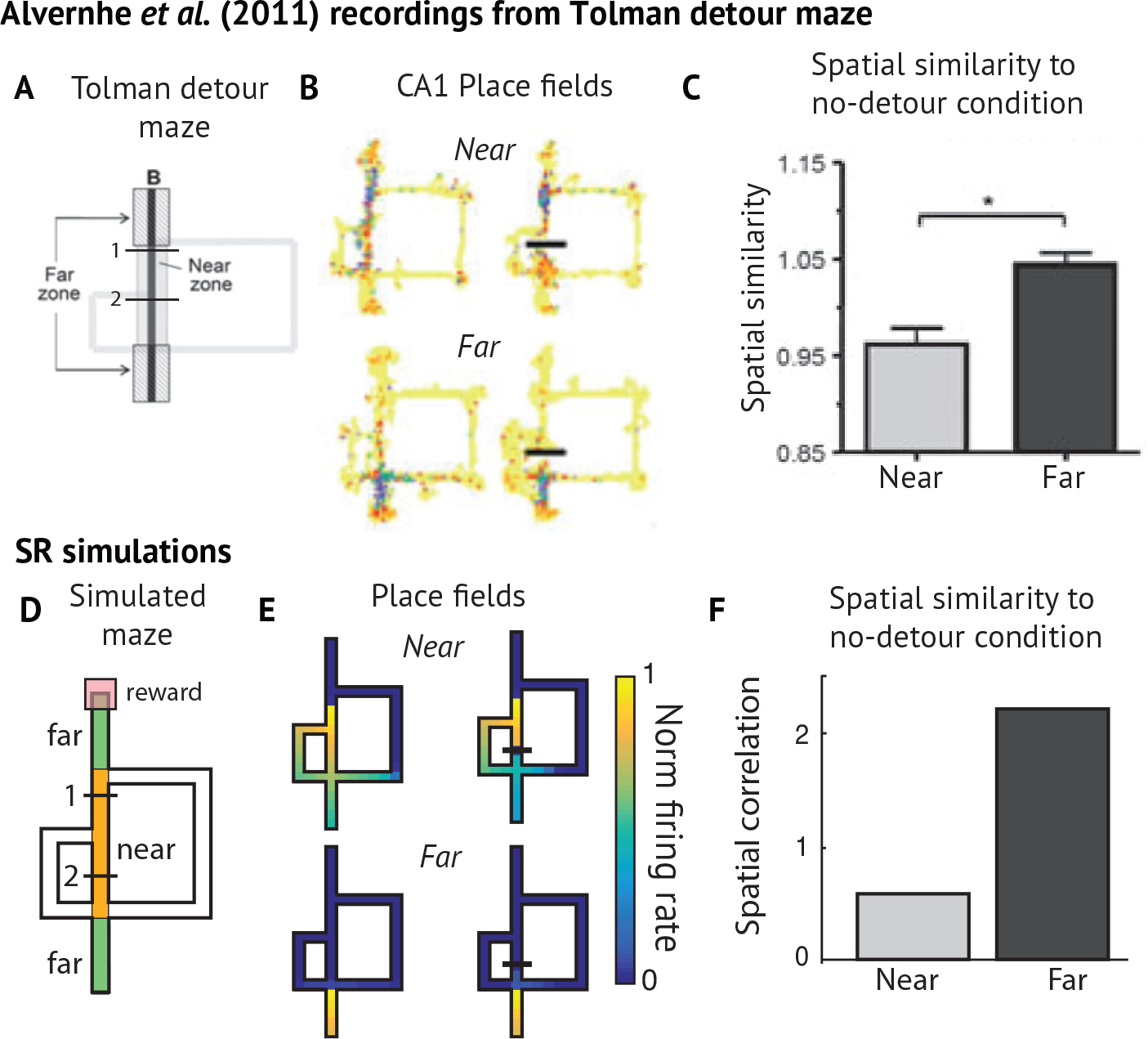
Place fields near detour. (A) Maze used by Alvernhe and colleagues^29^ for studying how place cell firing is affected by the insertion of barriers in a Tolman detour maze. Reward is delivered at location B. “Near” and “Far” zones are defined. In “early” and “late” detour conditions, a clear barrier blocks the shortest path, forcing the animal to take the short detour to the left or the longer detour to the right. (B) Example CA1 place fields recorded from a rat navigating the maze. (C) Over the population, place fields near the barrier changed their shape, while the rest remained unperturbed. This is shown by computing the Fisher *z* transformed spatial correlation between place field activity maps with and without barriers present. (D) The environment used to simulate the experimental results. (E) Example SR place fields near to and far from the barrier, before and after barrier insertion. More fields are shown in Fig. S6. (F) When barriers are inserted, SR place fields change their fields near the path blocked by the barrier and less so at more distal locations where policy is unaffected. The effect is more pronounced in the early detour condition because the detour appears closer to the start.

The SR model can be used to explain how hippocampal place fields depend on behaviorally relevant features that alter an animal's transition policy, such as reward. Using an annular watermaze, Hollup and colleagues demonstrated that a hidden, stationary reward affects the distribution of place fields^30^. Animals were required to swim in some preferred direction around a ring-shaped maze filled with an opaque liquid until they reached a hidden platform where they could rest. Hollup and colleagues found that the segment containing the platform had more place fields centered within it than any other segment, and that the preceding segment consistently had the second-largest number of place fields centered within it (Fig. 6A).

**Figure 6.**
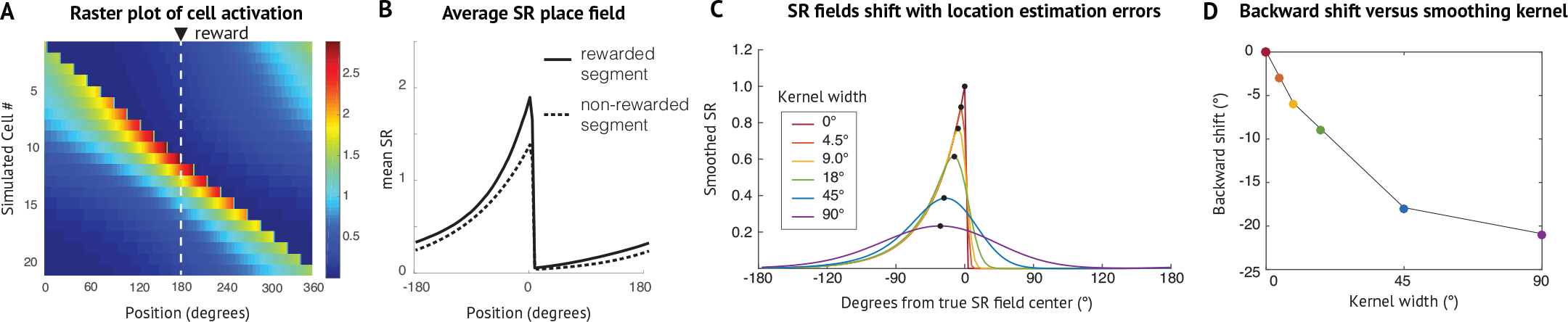
Distribution of place fields in annular maze with reward. (A) Simulated SR raster for annular watermaze. The transition model assumes that the animal spends more time near the rewarded platform and that the animal must move counter-clockwise (shown above as right-to-left) to get the reward. For this simulation, the probability of moving clockwise is 0. (B) The average SR place field in the rewarded and unrewarded segments. The states near the reward are visited more, so the SR model predicts more firing near these rewarded locations and the states that precede them. This difference is smaller when the discount factor is smaller. (C) When location is uncertain, the SR becomes smoother and the peak shifts toward the center of mass. For this reason, an asymmetric firing field may be accompanied by a backward migration of the firing field. (D) The magnitude of the shifts become more pronounced as the uncertainty distribution over possible locations of the animal becomes wider. For a given discount, the magnitude of the shift is bounded by distance between the SR field's center of mass and the encoded state.

We simulated this task using a sequence of states connected in a ring. The transition policy was such that the animal lingered longer near the rewarded location and had a preferred direction of travel (right, or counterclockwise, in this case), matching behavioral predictions recorded by the authors^30^. We set the probability of transitioning left to 0 to illustrate the predictions of our model more clearly. As we show in Figure 6A-B, the SR model predicts elevated firing near the rewarded location and backward skewing of place fields. This creates an asymmetry, whereby the locations preceding the rewarded location will experience slightly higher firing rates as well. Furthermore, this asymmetric backward skew makes it likely that fields will overlap with the previous segment, not the upcoming segment. Figure 6C-D demonstrates how this backward skewing can equate to a backward shift in cell peak in the presence of noise or location uncertainty. This may explain the asymmetry found in the distribution of place field peaks about the rewarded segment.

While Hollup and colleagues found an asymmetric distribution of place cells about the rewarded segment, they also found that place fields were roughly the same size at reward locations as at other locations. In contrast, the SR predicts that fields should get larger near reward locations (Fig. 6B), with the magnitude of this effect modulated by the discount factor (Fig. S6). Thus, the SR is still an incomplete account of reward-dependent place fields.

Note that the SR model does not predict that place fields would be immediately affected by the introduction of a reward. Rather, the shape of the fields should change as the animal gradually adjusts its policy and experiences multiple transitions consistent with that policy. The SR is affected by the presence of the reward because rewards induce a change in the animal's policy, which determines the predictive relationships between states.

Under a sufficiently large discount, the SR model predicts that firing fields centered near rewarded locations will expand to include the surrounding locations and increase their firing rate under the optimal policy. The animal is likely to spend time in the vicinity of the reward, meaning that states with or near reward are likely to be common successors. SR place fields in and near the rewarded zone will cluster because it is likely that states near the reward were anticipated by other states near the reward (Fig. S7). For place fields centered further from the reward, the model predicts that fields will skew opposite the direction of travel toward the reward, due to the effect illustrated in Fig. 2: a state will only be predicted when the animal is approaching reward from some more distant state. Given a large potentially rewarded zone or a noisy policy, these somewhat contradictory effects are sufficient to produce clustering of place fields near the rewarded zone (Fig. S7). The punished locations will induce the opposite effect, causing fields near the punished location to spread away from the rarely-visited punished locations (Fig. S5F). The SR place fields for each of these environments are shown in Figure S5.

In addition to the influence of experimental factors, changes in parameters of the model will have systematic effects on the structure of SR place fields. Motivated by data showing a gradient of increasing field sizes along the hippocampal longitudinal axis^31,32^, we explored the consequences of modifying the discount factor *γ* in Figure S4 and Figure S6. Hosting a range of discount factors along the hippocampal longitudinal axis provides a multi-timescale representation of space. It also circumvents the problem of having to assume the same discount parameter for each problem or adaptively computing a new discount. Another consequence is that larger place fields reflect the community structure of the environment. In Figure S5, we show how the SR fields begin to expand their fields to cover all states with the same compartment for a large enough discount. This overlap drives the clustering of states within the same community. A gradient of discount factors might therefore be useful for decision making at multiple levels of temporal abstraction^18,33,34^.

An appealing property of the SR model is that it can be applied to non-spatial state spaces. Fig. 7A-D shows the SR embedding of an abstract state space used in a study by Schapiro and colleagues^18,35^. Human subjects viewed sequences of fractals drawn from random walks on the graph while brain activity was measured using fMRI. We compared the similarity between SR vectors for pairs of states with pattern similarity in the hippocampus. The key experimental finding was that hippocampal pattern similarity mirrored the community structure of the graph: states with similar successors were represented similarly^35^. The SR model recapitulates these findings, since states in the same community tend to be visited nearby in time, making them predictive of one another (Fig. 7E-G). A recent related fMRI result from Garvert and colleagues provides further support that the hippocampus represents upcoming successors in a non-spatial, relational task by showing that a successor model provided the best metric for explaining variance in recorded hippocampal adaptation^36^.

**Figure 7.**
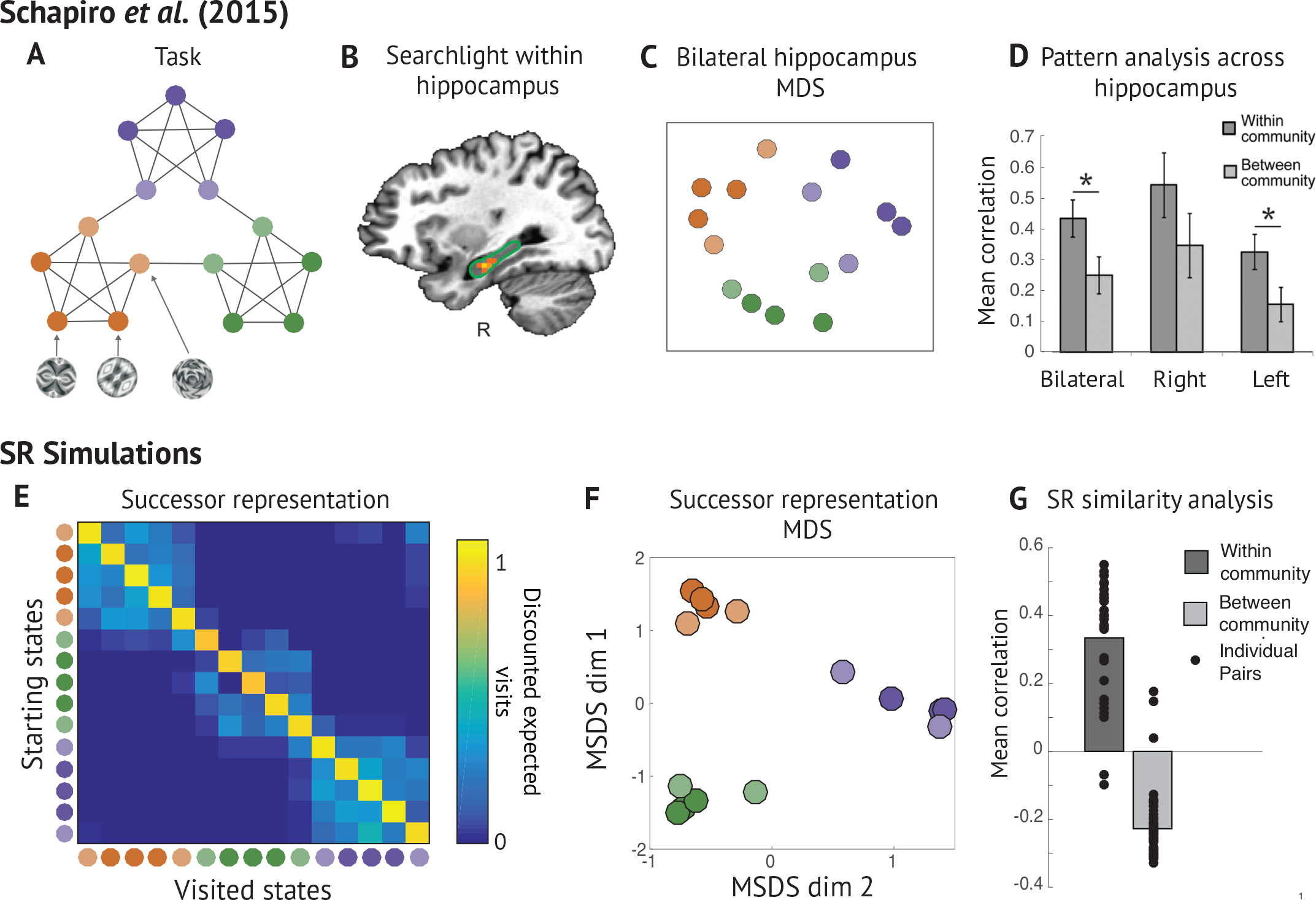
Hippocampal representations in non-spatial task. (A) Schapiro *et al*.^35^ showed subjects sequences of fractal stimuli drawn from the task graph shown, which has clusters of interconnected nodes (or “communities”). Nodes of the same color fall within the same community, with the lighter colored nodes connecting to adjacent communities. (B) A searchlight within hippocampus showed a stronger within-community similarity effect in anterior hippocampus. (C, D) States within the same cluster had a higher degree of representational similarity in hippocampus, and multidimensional scaling (MDS) of the hippocampal BOLD dissimilarity matrix captured the community structure of the task graph^35^. (E) The SR matrix learned on the task. The block diagonal structure means that states in the same cluster predict each other with higher probability. (F) Multidimensional scaling of dissimilarity between rows of the SR matrix reveals the community structure of the task graph. (G) Consistent with this, the average within-community SR state similarity is consistently higher than the average between-community SR state similarity.

To demonstrate further how the SR model can integrate spatial and temporal coding in the hippocampus, we simulated results from a recent study^37^ in which subjects were asked to navigate among pairs of locations to retrieve associated objects in a virtual city (8A). Since it was possible to “teleport” between certain location pairs, while others were joined only by long, winding paths, spatial Euclidean distance was decoupled from travel time. The authors found that objects associated with locations that were nearby in either space or time increased their hippocampal pattern similarity (Fig. 8B). Both factors (spatial and temporal distance) had a significant effect when the other was regressed out (Fig. 8C). The SR predicts this integrated representation of spatiotemporal distance: when a short path is introduced between distant states, such as by a teleportation hub, those states come predict one another.

**Figure 8.**
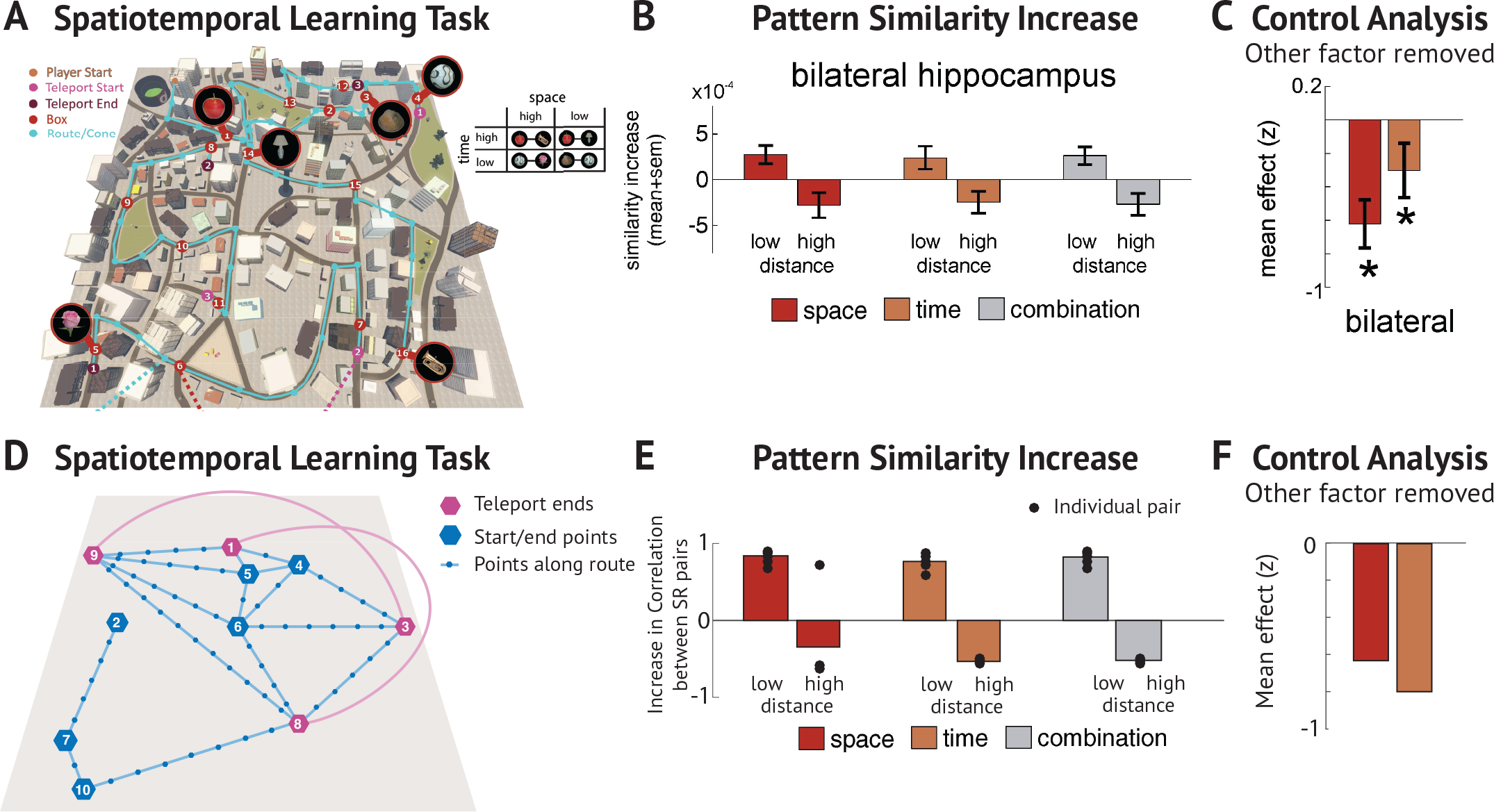
Hippocampal representations in spatio-temporal task. (A) Deuker *et al*.^37^ trained subjects on a spatio-temporal navigation task. Subjects were told to objects scattered about the map. It is possible to take a “teleportation” shortcut between certain pairs of states (pink and purple), and other pairs of states are sometimes joined only by a long, winding path. Nearness in time is therefore partially decoupled from nearness in space. (B) The authors find significant increase in hippocampal representational similarity between nearby states and a decrease for distant states. This effect holds when states are nearby in space, time, or both. (C) Since spatial and temporal proximity are correlated, the authors controlled for the each factor and measured the effect of the remaining factor on the residual. (D-F) Simulation of experimental results in panels A-C.

### Dimensionality reduction of the predictive map by entorhinal grid cells

Because the firing fields of entorhinal grid cells are spatially periodic, it was originally hypothesized that grid cells might represent a Euclidean spatial metric to enable dead reckoning^8,14^. Other theories have suggested that these firing patterns might arise from a low-dimensional embedding of the hippocampal map^5,23,38^. Combining this idea with the SR hypothesis, we argue that grid fields reflect a low-dimensional eigendecomposition of the SR. A key implication of this hypothesis is that grid cells will respond differently in environments with different boundary conditions.

The boundary sensitivity of grid cells was recently highlighted by a study that manipulated boundary geometry^39^. In square environments, different grid modules had the same alignment of the grid relative to the boundaries (modulo 60°, likely due to hexagonal symmetry in grid fields), whereas in a circular environment grid field alignment was more variable, with a qualitatively different pattern of alignment (Fig. 9A-C). Krupic *et al*. performed a “split-halves” analysis, in which they compared grid fields in square versus trapezoidal mazes, to examine the effect of breaking an axis of symmetry in the environment (Fig 9D,E). They found that moving the animal to a trapezoidal environment, in which the left and right half of the environment had asymmetric boundaries, caused the grid parameters to be different on the two sides of the environment^39^. In particular, the spatial autocorrelegrams – which reveal the layout of spatial displacement at which the grid field repeats itself – were relatively dissimilar over both halves of the trapezoidal environment. The grid fields in the trapezoid could not be attributed to linearly warping the square grid field into a trapezoid, raising the question of how else boundaries could interact with grid fields.

**Figure 9.**
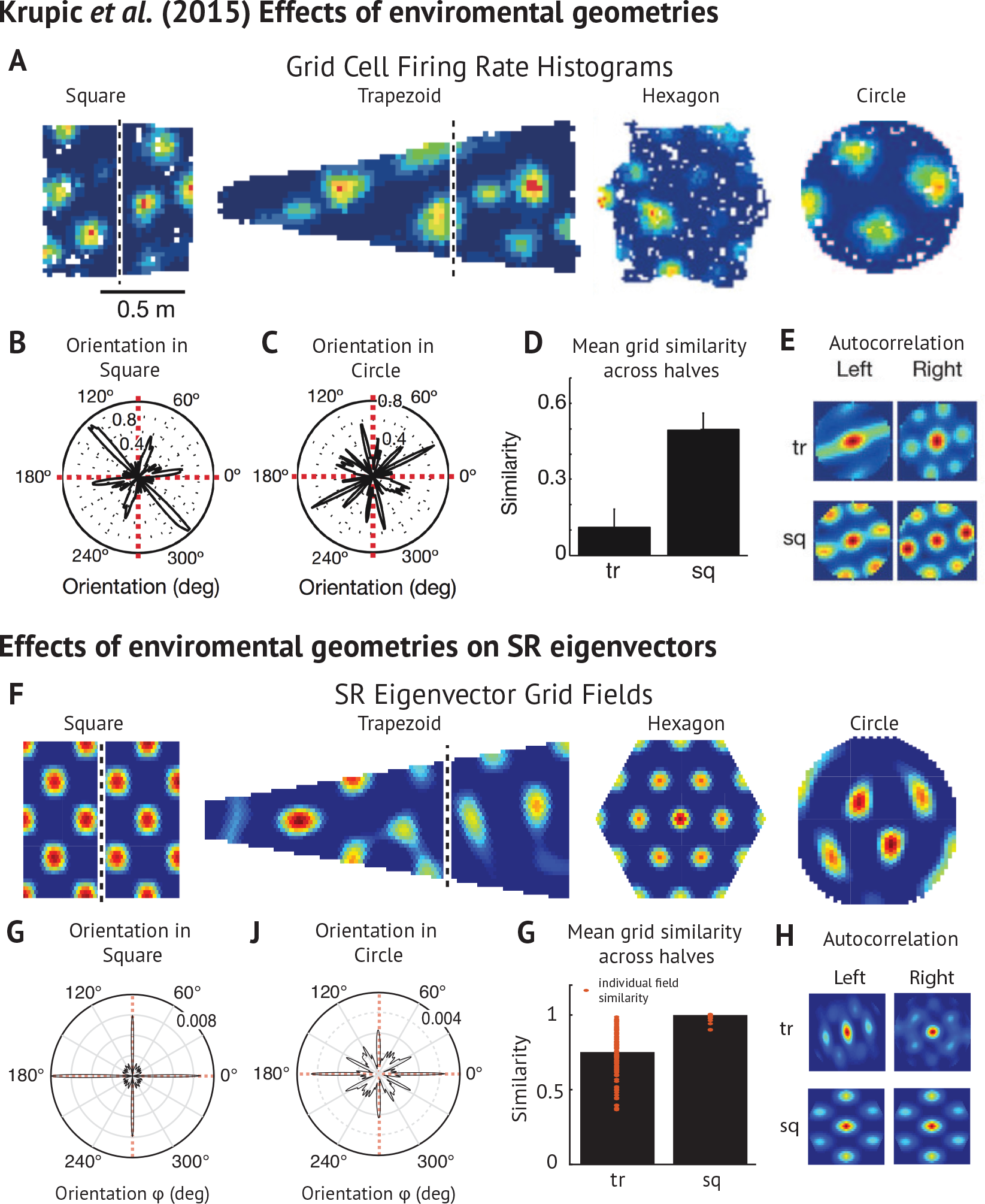
Grid fields in geometric environments. (A) Grid fields recorded in a variety of geometric environments^39^. Grid fields in trapezoid and square environments are split at the dividing line shown for split-halves analysis. (B,C) Grid fields in the square environment had more consistent orientations with respect to boundaries and distal cues than in the square environment. (D) While grid fields tend to be similar on both halves of a square (sq) environment, they tend to be less similar across halves of the irregular trapezoidal (tr) environment. (E) Autocorrelograms for different halves of trapezoidal and square environments in circular windows used for split-halves anal. (F-H) Simulations of experimental results in panels A-E.

According to the SR eigenvector model, these effects arise because the underlying statistics of the transition policy changes with the geometry. We simulated grid fields in a variety of geometric environments used by Krupic and colleagues (Fig. 9F-H; Fig. 9A–S9). In agreement with the empirical results, the orientation of eigenvectors in the circular environment tend to be highly variable, while those recorded in square environments are almost always aligned to either the horizontal or vertical boundary of the square (Fig. 9G,J). The variability in the circular environment arises because the eigenvectors are subject to the rotational symmetry of the circular task space. SR eigenvectors also emulate the finding that grids on either side of a square maze are more similar than those on either side of a trapezoid, because the eigenvectors capture the effect of these irregular boundary conditions on transition dynamics.

Another main finding of Krupic et al.^39^ was that when a square environment is rotated, grids remain aligned to the boundaries as opposed to distal cues. SR eigenvectors inherently reproduce this effect, since a core assumption of the theory is that grid firing is anchored to state in a transition structure, which is itself constrained by boundaries. The complete set of the first 64 eigenvectors is shown in Figures S8A and S9. While many fields conform to the canonical grid cell, others have skewed or otherwise irregular waveforms. Our model predicts that such shapes would be included in the greater variety of firing fields found in MEC that do not match the standard grid-like criterion.

A different manifestation of boundary effects is the fragmentation of grid fields in a hairpin maze^40^. Consistent with the empirical data, SR eigenvector fields tend to align with the arms of the maze, and frequently repeat across alternating arms (Figure 10)^40^. While patterns at many timescales can be found in the eigenvector population, those at alternating intervals are most common and therefore replicate the checkerboard pattern observed in the experimental data (Fig. S9).

**Figure 10.**
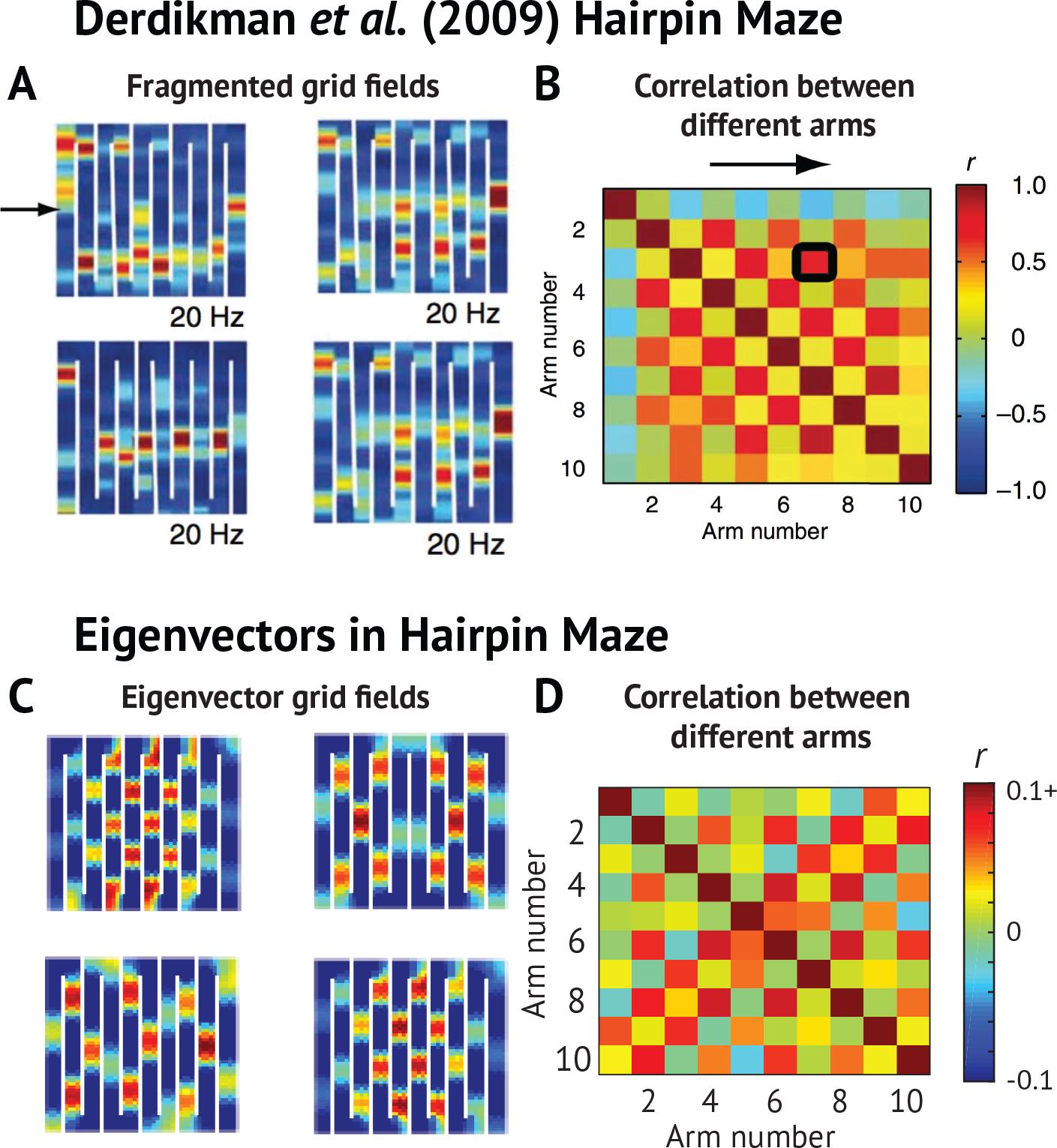
Grid fragmentation in compartmentalized maze. (A) Barriers in the hairpin maze cause grid fields to fragment repetitively across arms^40^. (B) Spatial correlation between activity in different arms. The checkerboard pattern emerges because grid fields frequently repeat themselves in alternating arms. (C-D) Simulations of the experimental results in panels A-B.

To further explore how compartmentalized environments could affect grid fields, we simulated a recent study^41^ that characterized how grid fields evolve over several days' exposure to a multi-compartment environment (Fig. 11). While grid cells initially represented separate compartments with identical fields (repeated grids), several days of exploration caused fields to converge on a more globally coherent grid (Fig. 11D-F). With more experience, the grid regularity of the fields simultaneously decreased, as did the similarity between the grid fields recorded in the two rooms (Fig. 11C). The authors conclude that grid cells will tend to a regular, globally coherent grid to serve as a Euclidean metric over the full expanse of the enclosure.

**Figure 11.**
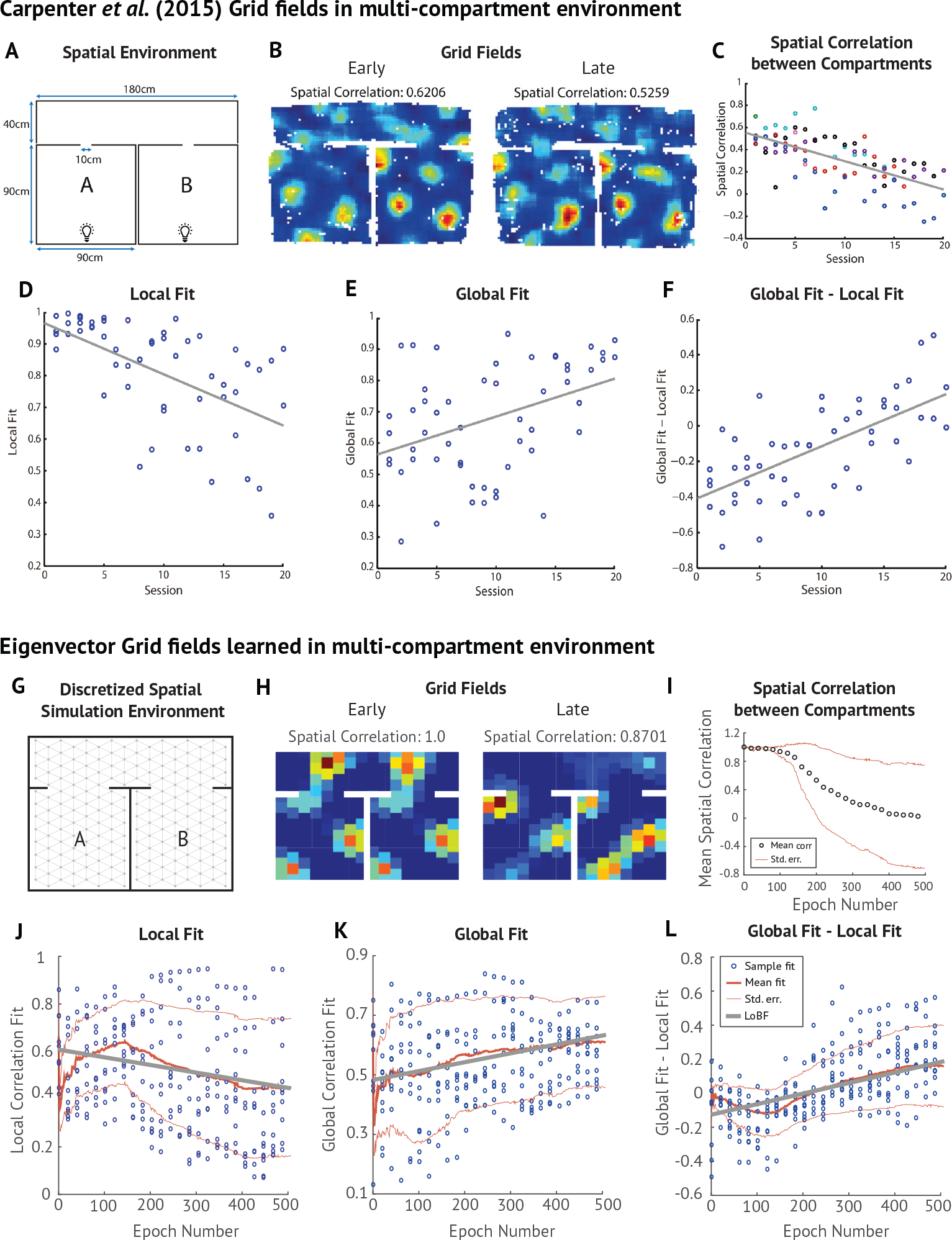
Grid fields in multi-compartment environment. (A) Multi-compartment environment employed by Carpenter and colleagues^41^. (B) Example grid fields early and late in training. (C) Spatial correlation between grid fields in compartments A and B across sessions. (D-F) To explain this decline in inter-compartment similarity, Carpenter and colleagues fit a local model (grid constrained to replicate between the two compartments) and a global model (single continuous grid spanning both compartments). They found that the local fit decreased across sessions, while the global fit increased, and correspondingly the difference between the two models increased. (G-L) Simulation of experimental results in panels A-F. In I-J, the blue circles indicate individual samples, the thick red line denotes the mean, the thin red lines denote one standard deviation from the mean, and the thick gray lines are lines of best fit.

Our model suggests that the fields are tending not toward a globally *regular* grid, but to a predictive map of the task structure, which is shaped in part by the global boundaries but also by the multi-compartment structure. We simulated this experiment by initializing grid fields to a local eigenvector model, in which the animal has not yet learned how the compartments fit together. After the SR eigenvectors have been learned, we relax the constraint that representations be the same in both rooms and let eigenvectors and the SR be learned for the full environment. As the learned eigenvectors converge, they increasingly resemble a global grid and decreasingly match the predictions of the local fit (Fig. 11H-L; Fig. S10). As with the recorded grid cells, the similarity of the fields in the two rooms drops to an average value near zero (Fig. 11I). They also have less regular grids compared to a single-compartment rectangular enclosure, explaining the drop in grid regularity observed by Carpenter *et al*. as the grid fields became more “global”^41^. Since separating barriers between compartments perturb the task topology from an uninterrupted 2D grid.

The eigenvectors of the SR are invariant to the discount factor of an SR matrix. This is because any SR can be written as a weighted sum of transition policy matrices, as we explain in more detail in the Supplemental Methods. The same eigenvectors will therefore support multiple SR matrices learned for the same task but with different planning horizons. SR matrices with a large discount factor will place higher eigenvalues on the eigenvectors with large spatial scales and low spatial frequency, whereas those with smaller discounts and smaller place fields project more strongly onto higher spatial-frequency grid fields. As discount is increased, the eigenvalues gradually shift their weight from the smaller scale to the larger scale eigenvectors (Fig. S11). This mirrors data suggesting that hippocampal connections to and from MEC vary gradually alongside place field spatial scale along the longitudinal axis^31, 32, 42, 43^. Grid fields, in contrast, cluster in discrete modules^44^. The SR eigenvectors are quantized as discrete modules as well, as we show in Figure S12.

A normative motivation for invoking low-dimensional projections as a principle for grid cells is that they can be used to smooth or “regularize” noisy updates of the SR. When the projection is based on an eigendecomposition, this constitutes a form of *spectral regularization*^45^. A smoothed version of the SR can be obtained by reconstructing the SR from its eigendecomposition using only low-frequency (high eigenvalue) components, thereby filtering out high-frequency noise (see Methods). This smoothing will fill in the blanks in the successor representations, enabling faster convergence time and a better approximation of the SR while it is still being learned. Spectral regularization has a long history of improving the approximation of large, incomplete matrices in real-world domains, most commonly through matrix factorization^45^. The utility of a spectral basis for approximating value functions in spatial and other environments has been demonstrated in the computational RL literature^46^. In Figure S13A, we provide a demonstration of how this kind of spectral regularization can allow the SR to be more accurately estimated despite the presence of corrupting noise in a multi-compartment environment. In Figure S13B, we show that spectral regularization provides a better reconstruction basis than a globally uniform Fourier basis, because the former does not smooth over boundaries.

We also demonstrate how reweighting eigenvalues so that more weight is placed on the low-frequency eigenvectors allows us to approximate the SR matrix for larger discounts with significantly less training time (Fig. S13C). TD learning can take a long time to converge when the discount factor is large. Spectral regularization can allow the SR to support planning over a longer timescale after significantly less training.

We include our own modest demonstration of how spectral regularization can improve SR-based value function approximation in a noisy, multicompartment spatial task. Importantly, the regularization is topologically sensitive, meaning that smoothing respects boundaries of the environment. Regularization using a Fourier decomposition does not share this property, and will smooth over boundaries (Fig. S13). The regularization hypothesis is consistent with data suggesting that although grid cell input is not required for the emergence of place fields, place field stability and organization depends crucially on input from grid cells^47–49^. These eigenvectors also provide a useful partitioning of the task space, as discussed in the following section.

### Subgoal discovery using grid fields

In structured environments, planning can be made more efficient by decomposing the task into subgoals, but the discovery of good subgoals is an open problem. The SR eigenvectors can be used for subgoal discovery by identifying “bottleneck states” that bridge large, relatively isolated clusters of states, and group together states that fall on opposite sides of the bottlenecks^50,51^. Since these bottleneck states are likely to be traversed along many optimal trajectories, they are frequently convenient waypoints to visit. Navigational strategies that exploit bottleneck states as subgoals have been observed in human navigation^52^. It is also worth noting that accompanying the neural results displayed in Fig. 7, the authors found that when subjects were asked to parse sequences of stimuli into events, stimuli found at topological bottlenecks were frequent breakpoints^18^.

The formal problem of identifying these bottlenecks is known as the *k*-way normalized min-cut problem. An approximate solution can be obtained using spectral graph theory^53^. First, the top log *k* eigenvectors of a matrix known as the graph Laplacian are thresholded such that negative elements of each eigenvector go to zero and positive elements go to one. Edges that connect between these two labeled groups of states are “cut” by the partition, and nodes adjacent to these edges are a kind of bottleneck subgoal. The first subgoals that emerge will be the cut from the lowest-frequency eigenvector, and these subgoals will approximately lie between the two largest, most separable clusters in the partition (see Supplemental Methods for more detail). A prioritized sequence of subgoals is obtained by incorporating increasingly higher frequency eigenvectors that produce partition points nearer to the agent.

The SR shares its eigenvectors with the graph Laplacian (see Supplemental Methods)^5^, making SR eigenvectors equally suitable for this process of subgoal discovery. We show in Figure S14 that the subgoals that emerge in a 2-step decision task and in a multi-compartment environment tend to fall near doorways and decision points: natural subgoals for high-level planning. It is worth noting that SR matrices parameterized by larger discount factors *γ* will project predominantly on the large-spatial-scale grid components (Fig. S11). The relationship between more temporally diffuse, abstract SRs, in which states in the same room are all encoded similarly (Fig. S4), and the subgoals that join those clusters can therefore be captured by which eigenvalues are large enough to consider.

It has also been shown experimentally that entorhinal lesions impair performance on navigation tasks and disrupt the temporal ordering of sequential activations in hippocampus while leaving performance on location recognition tasks intact^48,54^. This suggests a role of grid cells in spatial planning, and encourages us to speculate about a more general role for grid cells in hierarchical planning.

## Discussion

The hippocampus has long been thought to encode a cognitive map, but the precise nature of this map is elusive. The traditional view that the map is essentially spatial^7,8^ is not sufficient to explain some of the most striking aspects of hippocampal representation, such as the dependence of place fields on an animal's behavioral policy and the environment's topology. We argue instead that the map is essentially *predictive*, encoding expectations about an animal's future state. This view resonates with earlier ideas about the predictive function of the hippocampus^20, 24, 55–59^. Our main contribution is a formalization of this predictive function in a reinforcement learning framework, offering a new perspective on how the hippocampus supports adaptive behavior.

Our theory is connected to earlier work by Gustafson and Daw^13^ showing how topologically-sensitive spatial representations recapitulate many aspects of place cells and grid cells that are difficult to reconcile with a purely Euclidean representation of space. They also showed how encoding topological structure greatly aids reinforcement learning in complex spatial environments. Earlier work by Foster and colleagues^12^ also used place cells as features for RL, although the spatial representation did not explicitly encode topological structure. While these theoretical precedents highlight the importance of spatial representation, they leave open the deeper question of why particular representations are better than others. We showed that the SR naturally encodes topological structure in a format that enables efficient RL.

The work is also related to work done by Dordek *et al*.^23^, who demonstrated that gridlike activity patterns from principal components of the population activity of simulated Gaussian place cells. As we mentioned in the Results, one point of departure between empirically observed grid cell data and SR eigenvector account is that in rectangular environments, SR eigenvector grid fields can have different spatial scales aligned to the horizontal and vertical axis (see Fig. S8)^14^. In grid cells, the spatial scales tend to be approximately constant in all directions unless the environment changes^60^. The principal components of Gaussian place fields are mathematically related to the SR eigenvectors, and naturally also have grid fields that scale independently along the perpendicular boundaries of a rectangular room. However, Dordek *et al*. found that when the components were constrained to have non-negative values and the constraint that components be orthogonal was relaxed, the scaling became uniform in all directions and the lattices became more hexagonal^23^. This suggests that the difference between SR eigenvectors and recorded grid cells is not fundamental to the idea that grid cells are doing spectral dimensionality reduction. Rather, additional constraints such as non-negativity are required.

The SR can be viewed as occupying a middle ground between model-free and model-based learning. Model-free learning requires storing a look-up table of cached values estimated from the reward history^1,61^. Should the reward structure of the environment change, the entire look-up table must be re-estimated. By decomposing the value function into a predictive representation and a reward representation, the SR allows an agent to flexibly recompute values when rewards change, without sacrificing the computational efficiency of model-free methods^4^. Model-based learning is robust to changes in the reward structure, but requires inefficient algorithms like tree search to compute values^1,15^.

Certain behaviors often attributed to a model-based system can be explained by a model in which predictions based on state dynamics and the reward function are learned separately. For instance, the *context preexposure facilitation effect* refers to the finding that contextual fear conditioning is acquired more rapidly if the animal has the chance to explore the environment for several minutes before the first shock^62^. The facilitation effect is classically believed to arise from the development of a conjunctive representation of the context in the hippocampus, though areas outside the hippocampus may also develop a conjunctive representation in the absence of the hippocampus, albeit less efficiently^63^. The SR provides a somewhat different interpretation: over the course of preexposure, the hippocampus develops a *predictive* representation of the context, such that subsequent learning is rapidly propagated across space. Figure S15 shows a simulation of this process and how it accounts for the facilitation effect.

Many models of prospective coding in the hippocampus have drawn inspiration from the well-documented ordered temporal structure of firing in hippocampus relative to the theta phase^20,65,64^, and considered the many ways in which replaying hippocampal sweeps during sharp wave ripple events might be used for planning^66–71^. The firing of cells in hippocampus is aligned to theta such that cells encoding more distant places fire later during a theta cycle than immediately upcoming states (a phenomenon referred to as theta precession). States fire in a sequence ordered according to when they will next appear, suggesting a likely mechanism for forward sequential planning^65,72^.

However, precession alone is probably not sufficient to enact backward expansion of place fields in CA1, since NMDA antagonists that disrupt the persistent, backward expansion of place fields leave theta precession intact^73^. Furthermore, precession in CA1 likely originates outside of the hippocampus, as it arises in MEC independently^75^, and depends crucially on input from surrounding areas such as MEC and CA3^54,75^. Thus, we think that it is worthwhile to consider the possible contributions of this backward expansion to planning in addition to the contributions of the hippocampal temporal code examined by this prior work.

The type of prospective coding implemented by theta precession and sharp wave ripple events is reminiscent of model-based, sequential forward planning^20^; many experiments and theoretical proposals have looked at how replaying these sequences at decision points and at rest can underlie planning^66–68, 70, 71^. By integrating the reward reactivated at each state along a sweep through upcoming states, the value of a specific upcoming trajectory can be predicted.

The SR is a different type of prospective code, with different tradeoffs. The SR marginalizes over all possible sequences of actions, making predictions over an arbitrarily long timescale in constant time. This results in a loss of flexibility relative to model-based planning, but greater computational efficiency. Thus, the SR cannot replace the full functionality of model-based sweeps. However, it might provide a useful adjunct to this functionality.

One way to combine the strengths of model-based planning with the SR would be to use the SR to extend the range of forward sweeps. In Fig. S19, we illustrate how performing sweeps in the successor representation space (Fig. S19F) or performing sweeps that terminate on a successor representation of the terminal state (Fig. S19G) can extend the range of these predictions, making the hippocampal representations a more powerful substrate for planning. This is tantamount to a “bootstrapped search” algorithm, variants of which have been successful in a range of applications^76,77^.

The SR model we describe is trained on the policy the animal has experienced. This means that when the reward is changed, the new value function computed from the existing SR will initially be based on the old policy. The new optimal policy is unlikely to be the same as the old one, which means that the new value function is not correct, and must be refined as the animal optimizes its behavior. This problem is encountered with all learning algorithms that learn cached statistics under the current policy dynamics.

In some cases, the old SR will be a reasonable initialization. In many environments, certain aspects of the dynamics are not subject to the animal's control, and the underlying adjacency structure is unlikely to change. Furthermore, if rewards tend to be distributed in the same general area of a task, many policy components will generalize. It is hard to make comprehensive claims about whether or not the space of naturalistic tasks adheres to these properties in general. Recent computational work has demonstrated that deep successor features (a more powerful generalization of the successor representation model) generalize well across changing goals and environments in the domain of navigation^78^.

To give an intuition of how the flexibility of the SR-based value computation depends on task hierarchy and simulation parameters, we look at generalization using a simple tree-structured maze. Figure S16 illustrates how the quality of SR generalization depends on the policy stochasticity (parameterized by *β*) and how similar the optimal paths are for the old and new rewarded location. When there is greater stochasticity (closer to the random walk policy), the SR's generalization to highly dissimilar locations is less impaired, but there is also a reduced generalization advantage when the reward ends up nearby. The random walk SR is used as a baseline. By diffusing value through the graph in accordance with the task's underlying adjacency structure, this representation always generalizes better than re-initializing to a state index representation. The animal should maintain support for random actions until it is very certain of the optimal path. Spectral regularization can promote this by smoothing the SR.

When the SR fails to support value computation given the new reward, there are other mechanisms that can compensate. Models such as Dyna update cached statistics using sweeps through a model, revising them flexibly^76^. The original form of Dyna demonstrated how model-based and model-free mechanisms could collaboratively update a value function. However, the value function can be replaced with any statistic learnable through temporal differences, including the SR, as demonstrated by recent work^79^. Furthermore, there is evidence from humans that when reward is changed, revaluation occurs in a policy-dependent manner, consistent with the kind of partial flexibility conferred by the SR^80^.

Recent work has elucidated connections between models of episodic memory and the SR. Specifically, Gershman *et al*. demonstrated that the SR is closely related to the Temporal Context Model (TCM) of episodic memory^16,19^. The core idea of TCM is that items are bound to their temporal context (a running average of recently experienced items), and the currently active temporal context is used to cue retrieval of other items, which in turn cause their temporal context to be retrieved. The SR can be seen as encoding a set of item-context associations. The connection to episodic memory is especially interesting given the crucial mnemonic role played by the hippocampus and entorhinal cortex in episodic memory. Howard and colleagues^81^ have laid out a detailed mapping between TCM and the medial temporal lobe (including entorhinal and hippocampal regions).

Spectral graph theory provides insight into the topological structure encoded by the SR. We showed specifically that eigenvectors of the SR can be used to discover a hierarchical decomposition of the environment for use in hierarchical RL. Spectral analysis has also frequently been invoked as a computational motivation for entorhinal grid cells (e.g.,^82^). The fact that any function can be reconstructed by sums of sinusoids suggests that the entorhinal cortex implements a kind of Fourier transform of space.

However, Fourier analysis is not the right mathematical tool when dealing with spatial representations in a topologically structured environment, since we do not expect functions to be smooth over boundaries in the environment. This is precisely the purpose of spectral graph theory: Instead of being maximally smooth over Euclidean space, the eigenvectors of the graph Laplacian embed the smoothest approximation of a function that respects the graph topology^46^.

In conclusion, the SR provides a unifying framework for a wide range of observations about the hippocampus and entorhinal cortex. The multifaceted functions of these brain regions can be understood as serving a superordinate goal of prediction.

## Methods

### Task simulation

Environments were simulated by discretizing the plane into points, and connecting these points along a triangular lattice (Fig. S1A). The adjacency matrix *A* was constructed such that *A*(*s, s*′) = 1 wherever it is possible to transition between states *s* and *s*′, and 0 otherwise.

The transition probability matrix *T* was defined such that *T* (*s, s*′) is the probability of transitioning from state *s* to *s*′. Under a random walk policy, where the agent chooses randomly among all available transitions, the transition probability distribution is uniform over allowable transitions. This amounts to simply normalizing *A* so that each row of *A* sums to 1 to meet the constraint that the possible transition from *s* must sum to 1. When reward or punishment was included as part of the simulated task, we computed the optimal policy using value iteration and a softmax value function parameterized by *β* ^15^.

### SR computation

The successor representation is a matrix, *M* where *M*(*s, s*′) is equal to the discounted expected number of times the agent visits state *s*′ starting from *s* (see Equation 3 for the mathematical definition and Fig. S1B for an illustration). When the transition probability matrix is known, we can compute the SR as a discounted sum over transition matrices raised to the exponent *t*. The matrix *T^t^* is the *t*-step transition matrix, where *T^t^* (*s, s*′) is the probability of transitioning from *s* to *s*′in exactly *t* steps.

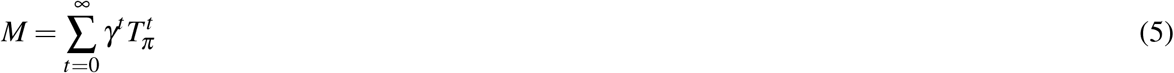

This sum is a geometric matrix series, and for *γ* < 1, it converges to the following finite analytical solution:

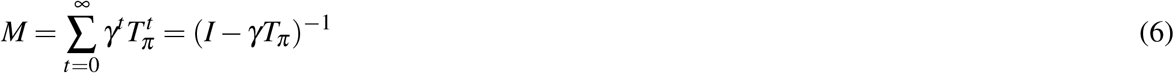

In most of our simulations, the SR was computed analytically from the transition matrix using this expression.

The SR can be learned on-line using the temporal differences update rule of Equation 4^4^ (also see^15^ for background on TD learning) (Fig. 11, Fig. S1, Fig. S3). When noise was injected into the location signal (Fig. S3). Noise was injected into the location signal by adding uniform random noise with mean 0 to the state indicator vector.

### Eigenvector computation and Spectral Regularization

In generating the grid cells shown, we assume random walk policy, which is the maximum entropy prior for policies (see^84^ for why maximum entropy priors can be good priors for regularization). However, since the learned eigenvectors are sensitive to the sampling statistics, our model predicts that regions of the task space more frequently visited would come to be over-represented in the grid space (see Figure S8 for examples). For most figures, we compute the eigenvectors of the SR using the built-in MATLAB eig function (Fig. S1C). We then thresholded the eigenvectors at 0 so that firing rates are not negative (Fig. S1D).

For Figure 11, eigenvectors were computed incrementally using a Candid Covariance-free Incremental PCA (CCIPCA), an algorithm that efficiently implements stochastic gradient descent to compute principal components^83^ (eigenvectors and principal components are equivalent in this and many domains). Spectral regularization was implemented by reconstructing the SR from the truncated eigendecomposition (Fig. S13). Spectral reconstruction for Figure S13 was implemented by shifting the eigenvalues so that more weight was placed on low-frequency eigenvectors, rather than imposing a hard cutoff on high-frequency eigenvectors, and reconstructing an SR that corresponded to a larger discount factor. This allowed larger-discount SRs to be more exactly approximated. The reconstructed SR matrices *M*_recon_ were compared to the ground truth matrix *M*_gt_ by taking the correlation between *M*_recon_ and *M*_gt_ (Fig. S13). This measure indicates whether policies based on SR-based value functions for different reward functions will to tend send the animal in the right direction. Details can be found in the Supplemental Methods.

### Plotting receptive fields

To visualize place fields under the SR model, we showed heat maps of how active each SR-encoding neuron would be at each state in the environment (Fig. S1E-F). This shows the discounted expected number of times the neuron's encoded state *s* will be visited from each other state in the environment, and corresponds to taking a column *M*(*s*,:) from the SR matrix and reshaping it so that each element appears at the *x, y* location of its corresponding state. We use the same reshaping and plotting procedure to visualize eigenvector grid cells, using the columns of the thresholded eigenvector matrix *U* in place of *M*.

### Quantifying place and grid fields

To quantify place field clustering, center of mass (CoM) of SR place fields was computed by summing the locations of firing, weighted by the firing rate at that location (normalized so that the total firing summed to 1):

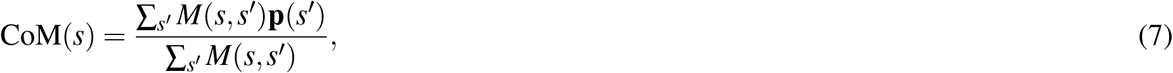

where **p**(*s*′) is the (*X,Y*) coordinate of the place field centered at state *s*′.

In Fig. 5, spatial similarity was computed by taking the Fisher *z* transform of spatial correlation between fields. Statistics were computed in this *z* space.

Grid field quantifications paralleled the analyses of Krupic *et al*.^39^: an ellipse was fit to the 6 peaks closest to the central peak, “orientation” refers to the orientation of the main axes (*a*,*b*). “Correlation” always refers to the Pearson correlation, “spatial correlation” refers to the Pearson correlation computed over points in space (as opposed to points in a vector), and spatial autocorrelation refers to the 2D auto-convolution.

To measure similarity between halves of the environment in Figure 9, we 1) computed the spatial autocorrelation for each half, 2) selected a circular window in the center of the autocorrelation, and 3) computed the correlation between autocorrelations of the two halves in the window. This paralleled the analysis taken by Krupic *et al*.^39^ and provides a measure of grid similarity across halves of the environment. The circular window is used to control for the fact that the boundaries of the square and trapezoid in the two halves of the respective environments differ. The mean similarity was *not* computed in Fisher *z*-transformed space, as one would normally do, but rather in correlation space. This was because the similarity for many of the square eigenvectors and at least one trapezoidal eigenvector was exactly 1, for which *z* = ∞. A dot plot is superimposed over this plot so the statistics of the distribution can be visualized.

In evaluating our simulations of the grid fields reported by Carpenter *et al*.^41^ (Fig. 11), the local model consisted of the set of 2D Fourier components bounded by the size of the compartment and the global model consisted of the set of 2D Fourier components bounded by the size of the environment. “Model fit” was measured for each eigenvector by finding maximum correlation over all model components between the eigenvector and model component.

#### Code availability

These results were generated using code written in MATLAB. If you are interested in accessing the code, you can email the corresponding author and we will be happy to make it available.

## Acknowledgments

We are grateful to Tim Behrens, Ida Mommenejad, and Kevin Miller for helpful discussions, and to Alexander Mathis and Honi Sanders for comments on an earlier draft of the paper. This research was supported by the NSF Collaborative Research in Computational Neuroscience (CRCNS) Program Grant IIS-120 7833 and The John Templeton Foundation. The opinions expressed in this publication are those of the authors and do not necessarily reflect the views of the funding agencies.

## Author contributions statement

All authors conceived the model and wrote the manuscript. Simulations were carried out by K.S.

## Additional information

The authors declare no competing financial interests.

